# Glycopeptide variable window SWATH for improved Data Independent Acquisition glycoprotein analysis

**DOI:** 10.1101/739615

**Authors:** Chun Zhou, Benjamin L. Schulz

## Abstract

*N*-glycosylation plays an essential role in regulating protein folding and function in eukaryotic cells. Sequential window acquisition of all theoretical fragment ion spectra mass spectrometry (SWATH) has proven useful as a data independent acquisition (DIA) MS method for analysis of glycoproteins and their glycan modifications. By separating the entire *m/z* range into consecutive isolation windows, DIA-MS allows comprehensive MS data acquisition and high-sensitivity detection of molecules of interest. Variable width DIA windows allow optimal analyte measurement, as peptide ions are not evenly distributed across the full *m/z* range. However, the *m/z* distribution of glycopeptides is different to that of unmodified peptides because of their large glycan structures. Here, we improved the performance of DIA glycoproteomics by using variable width windows optimized for glycopeptides. This method allocates narrow windows at *m/z* ranges rich in glycopeptides, improving analytical specificity and performance. We show that related glycoforms must fall in separate windows to allow accurate glycopeptide measurement. We demonstrate the utility of the method by comparing the cell wall glycoproteomes of wild-type and *N*-glycan biosynthesis deficient yeast and showing improved measurement of glycopeptides with different glycan structures. Our results highlight the importance of appropriately optimized DIA methods for measurement of post-translationally modified peptides.

## Introduction

Protein glycosylation is a common and complex co- and post-translational modification (PTM), which provides proteome diversity, regulates protein folding, sorting and stability, and controls many protein functions [1–7]. Different forms of glycosylation are defined by the linkages between glycans and proteins. *N*-glycosylation, occurring on specific asparagine (Asn) residues, is one of the most common forms of glycosylation in eukaryotic cells [8, 9]. *N*-glycans have a conserved precursor oligosaccharide, a dolichol-linked 14-monosaccharide molecule Glc_3_Man_9_GlcNAc_2_ [10–12]. This common core is assembled on the endoplasmic reticulum (ER) membrane, catalyzed by a series of glycosyltransferases encoded by asparagine-linked glycosylation (*ALG*) genes [13]. Oligosaccharyltransferase (OTase) catalyzes the attachment of the precursor oligosaccharide to selected Asn residues in glycosylation sequons (N-X-S/T; X≠P) in nascent polypeptides [14]. Mutations in *ALG* genes can cause altered glycan structures due to the accumulation and transfer of truncated glycans, and altered site-specific glycan occupancy due to inefficient transfer of these truncated glycans by OTase [15, 16]. Further trimming and maturation of glycans take place in the ER and Golgi with various enzymes involved, leading to diverse *N*-glycan structures on mature glycoproteins [17, 18].

*N*-glycosylation depends on both the genetics and environment of a cell, as the *N*-glycosylation pathway can be influenced by the physiological state of the cell, leading to changes in the sitespecific structures and heterogeneity of *N*-glycans across the glycoproteome [19–21]. Efficient and robust analysis of the heterogeneity of the glycoproteome is therefore critical [22]. In practice, glycoproteomic analyses have been used to better understand the molecular bases and consequences of various diseases [22–26], including congenital disorders of glycosylation (CDG), a family of diseases caused by defects in glycosylation biosynthesis pathways [27]. Glycoproteins and glycans also have potential as biomarkers for early detection, diagnosis, and prognosis of diseases including diverse cancers [19, 28–30].

Mature glycoproteins can have an incredible diversity in the site-specific presence and structures of glycans [31]. This high diversity and relatively low intrinsic detectability of glycosylated peptides further increases the challenges in glycosylation analysis in mass spectrometry (MS) glyco/proteomic workflows. Further improvements in sensitive, robust, high-throughput, and efficient methods for site-specific analysis of glycoprotein structure and occupancy are therefore required. MS-based techniques are the current method of choice for glycoproteomic analyses, as they offer high speed, resolution, and sensitivity [32, 33]. A wide range of MS strategies have been described for quantifying site-specific glycosylation in complex protein samples either in glycoproteomic workflows with glycans still attached to peptides [34–36], or in glycomic workflows that analyse glycans that have been released from glycoproteins [37], either labelled [38–40] or label-free [41, 42].

A variety of MS data acquisition approaches are available for proteomic studies, depending on the experimental design and analytical goal [43–45]. Data Independent Acquisition (DIA) methods such as sequential window acquisition of all theoretical fragment ion spectra mass spectrometry (SWATH) are especially useful for comprehensive measurement of all detectable analytes [46]. DIA/SWATH has been useful in analysis of complex proteomes in many diverse systems, including investigations of fundamental cell biology, biomarker discovery, and analysis of proteoform diversity [46–48]. Glycoproteomic analyses investigating the role of the glycosylation machinery in glycoprotein synthesis and identification of cancer biomarkers have also successfully used DIA/SWATH [34, 49, 50].

In a standard implementation of DIA/SWATH, the *m/z* field is broken into consecutive 25 *m/z* fixed width isolation windows (swaths) with a small overlap [46]. However, the distribution of peptides across the *m/z* range is uneven, leading to unbalanced precursor densities in each window, reducing analytical performance. The spectral complexity and chances of incorrect peak measurement in MS/MS spectra and extracted fragment ion chromatograms (XICs) are higher for the windows containing more precursor ions, while windows with few precursor ions do not make efficient use of MS instrument cycle time [51, 52]. These problems have been overcome by the use of variable window DIA/SWATH, an acquisition strategy that uses DIA isolation windows of different widths to equalize the average distribution of precursor ions in each window. This approach improves selectivity, detection confidence, and quantification reliability [51, 52]. Several strategies for generating variable windows for DIA/SWATH have been used to allow improved identification, profiling, and quantification of peptides and their modifications [52–55]. For example, equalizing either the precursor ion population or the total ion current in each window using the web application SwathTUNER increased the number of identified peptide precursors and proteins [52]. Additionally, separating the full mass range into more width-diverse windows improved the measurement of lysine acetylation and succinylation [54]. However, the high mass of glycopeptides means these previous approaches are not optimal for DIA/SWATH glycoproteomic analyses. This is especially problematic for DIA/SWATH measurement of glycopeptides that happen to fall in the same window, as differently glycosylated forms of the same underlying peptide share most fragment ions. Here, we describe a glycopeptide-focused variable window DIA/SWATH method that improves measurement of *N*-glycosylated peptides.

## Methods

### Yeast Strain and Growth Conditions

Yeast strains used in this study were BY4741 wild-type (*MATa his3*Δ*1 leu2*Δ*0 met15*Δ*0 ura3*Δ*0*) and the *alg3Δ* mutant (*MATa his3*Δ*1 leu2*Δ*0 met15*Δ*0 ura3*Δ*0 alg3*Δ*::kanMX*, Open Biosystems). YPD medium containing 1% yeast extract, 2% peptone and 2% glucose was used for yeast growth.

### Cell Wall Protein Sample Preparation

Cell wall proteins were enriched as previously described [34, 56]. In brief, yeast cells were grown in YPD at 30 °C with shaking until the OD_600nm_ reached 1.0. Cells were harvested by centrifugation and completely lysed by vortex with glass beads at 4 °C. Proteins were denatured, cysteines were reduced with 10 mM dithiothreitol and alkylated with 25 mM acrylamide. Proteins covalently linked to the yeast polysaccharide cell wall were isolated by centrifugation, and non-covalently linked contaminating proteins were removed by washing with strongly denaturing buffer containing 2% SDS, 7 M urea and 2 M thiourea. Proteins in the insoluble cell wall fraction were digested by trypsin and peptides and glycopeptides were desalted using C18 ZipTips (Millipore).

### Variable Window SWATH Method Generation

A variable window SWATH method designed for glycopeptide measurement (gpvwSWATH) was generated by the program SWATH variable window calculator Version 1.0 (SCIEX) using a previous yeast cell wall MS proteomics dataset [34]. Peptides with MS/MS spectra containing HexNAc and Hex oxonium ions were considered as glycopeptides [57]. Glycopeptides in the *m/z* range of 400-1250 were selected as input for generating the gpvwSWATH design. In addition, a traditional variable window SWATH (vwSWATH) method was generated using all peptides within the same *m/z* range. Both variable window SWATH methods had 34 windows over the *m/z* range of 400-1250 Da, consistent with the fixed window SWATH and the maximum *m/z* of the TripleTof 5600 instrument. Window overlap was 1 Da, and the minimum window width was set as 3 Da.

### Mass Spectrometry

Desalted cell wall peptides were analysed by LC-ESI-MS/MS using a Prominence nanoLC system (Shimadzu) and TripleTof 5600 instrument (SCIEX) as previously described [34, 37]. Approximately 2 μg peptides were desalted on an Agilent C18 trap (300 Å pore size, 5 μm particle size, 0.3 mm i.d. x 5 mm) at a flow rate of 30 μl/min for 3 min, and then separated on a Vydac EVEREST reversed-phase C18 HPLC column (300 Å pore size, 5 μm particle size, 150 μm i.d. x 150 mm) at a flow rate of 1 μl/min. Peptides were separated with a gradient of 10-60 % buffer B over 14 min, with buffer A (1 % acetonitrile and 0.1 % formic acid) and buffer B (80 % acetonitrile with 0.1 % formic acid). Gas and voltage setting were adjusted as required. An MS TOF scan from *m/z* of 350-1800 was performed for 0.5 s followed by data dependent acquisition (DDA) of MS/MS with automated CE selection of up to 20 peptides from *m/z* of 350-1250 for 0.05 s per spectrum. Proteins were detected and quantified using DIA/SWATH as previously described [34, 58], using identical LC parameters as for DDA analysis. For fixed window or variable window SWATH-MS, an MS-TOF scan was performed for 0.05 s, followed by MS/MS for 0.1 s for each isolation window.

### Data Analysis

A previously generated ion library was used for glycopeptide quantification [34]. This ion library contained eight possible cell wall glycopeptides, and included b and y fragment ions of non-glycosylated sequon-containing peptides and precursor ion masses of every possible glycopeptide with glycan structures ranging from GlcNAc_2_ to Man_15_GlcNAc_2_. Peptide measurement was performed with PeakView as previously described [34]. The mass spectrometry proteomics data have been deposited to the ProteomeXchange Consortium via the PRIDE [59] partner repository with the dataset identifier PXD015043. The abundance of each peptide was measured by summing the integrated areas of up to six fragment ions. Low-quality data was eliminated using a python script based on the co-generated FDR file as previously described [60], with only peptide intensity values with a corresponding FDR less than 0.01 accepted. The intensity of each peptide glycoform was normalized against the sum of all glycoforms of that peptide. Statistical analyses of peptide intensity differences between wild type and *alg3Δ* were performed using *t*-tests, with *p* < 0.05 considered to be significant. All experiments were performed in biological triplicate.

## Results

### Generating a glycopeptide focussed variable window SWATH method

The distribution of *m/z* values and intensities of precursor ions of the analysts observed from LC-MS/MS analysis of a comparable experimental sample are generally used to generate a variable window SWATH method. This data can be used as input to a variable window calculator [54, 55]. The SWATH variable window calculator allocates window widths to obtain a roughly equal density of precursor ions in each window. We generated two variable window SWATH methods based on input data from LC-MS/MS proteomic analysis of yeast cell wall peptides and glycopeptides from wild type and several *N*-glycosylation pathway deficient yeasts [34].

The first variable window SWATH method (vwSWATH) was optimized for all detectable peptides in the dataset [34]. Based on the traditional variable window SWATH generation approach, all MS1 peptide ions within the *m/z* range of 400 to 1250 Da were used (Fig. 1A). A pronounced uneven distribution of ions was observed with a peak density of precursor ions at ~550 *m/z* and the bulk of ions falling between 400-800 *m/z*. Splitting the full *m/z* range into 34 windows of variable width produced windows ranging from 13.5 to 87.3 *m/z* in width (Table 1). Most windows fell in the ion dense region between an *m/z* of 400 to 800, while windows outside of this region were much broader. This distribution is typical of LC-MS/MS peptide profiles.

**Figure 1.**
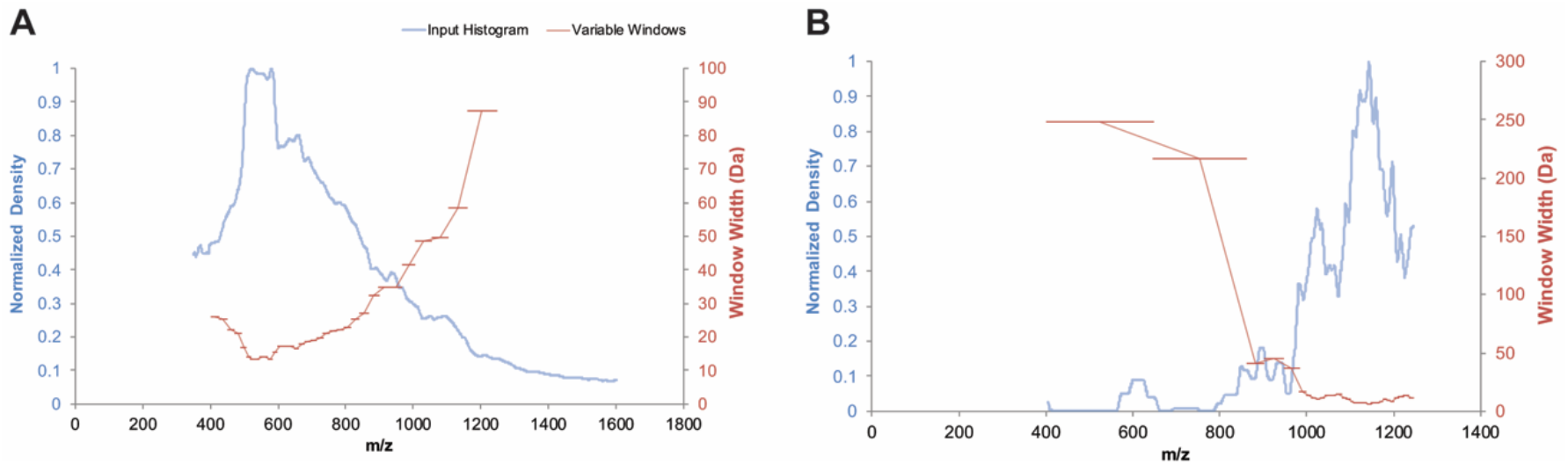
Design of variable window SWATH methods for peptides and glycopeptides. **(A)** Traditional peptide-focused variable window SWATH. Blue line, distribution of all peptide precursor ions. **(B)** Glycopeptide-focused variable window SWATH. Blue line, distribution of glycopeptide precursor ions. Red line, window width allocation. Each method had 34 windows.

Due to the presence of large and complex glycans in addition to the peptide backbone, glycopeptides are typically larger than peptides and have a different average *m/z* distribution. Variable window SWATH methods optimized for peptides are therefore unlikely to be optimal for measurement of glycopeptides. We therefore next designed a variable window SWATH method optimized for measurement of glycopeptides (gpvwSWATH). We used the same previous analysis used for vwSWATH generation [34], and created a list of *m/z* values of yeast cell wall glycopeptides in an unbiased and sensitive manner by selecting MS/MS spectra that contained glycan-specific oxonium fragment ions from HexNAc and Hex monosaccharides. Glycopeptide ions within the *m/z* range of 400 to 1250 Da were then used to generate the gpvwSWATH method, with 34 windows across the *m/z* range of 400 to 1250 Da (Fig.1B). The *m/z* distribution of glycopeptides was even more uneven than for peptides, leading to widely divergent window widths. As almost all glycopeptide precursor ions were larger than 800 *m/z*, windows at low *m/z* values were often wider than 200 Da (Table 1), while windows between 900-1200 *m/z* were narrow due to the high glycopeptide precursor ion density in this region. The shift of the precursor ion dense region from the centre of the mass range towards higher *m/z* values is consistent with attachment of glycans substantially increasing the average *m/z* distribution of glycopeptides.

**Table 1.** Window distributions of two variable window SWATH methods

| Variable Window SWATH<br>(vwSWATH) |  |  | GlycoPeptide Variable Window SWATH<br>(gpvwSWATH) |  |  |
| --- | --- | --- | --- | --- | --- |
| start $m/z$ | end $m/z$ | window width | start $m/z$ | end $m/z$ | window width |
| 399.5 | 425.5 | 26.0 | 399.5 | 647.7 | 248.2 |
| 424.5 | 449.9 | 25.4 | 646.7 | 863.3 | 216.6 |
| 448.9 | 471.1 | 22.2 | 862.3 | 903.2 | 40.9 |
| 470.1 | 491.1 | 21.0 | 902.2 | 947.7 | 45.5 |
| 490.1 | 506.8 | 16.7 | 946.7 | 983.7 | 37.0 |
| 505.8 | 519.9 | 14.1 | 982.7 | 999.3 | 16.6 |
| 518.9 | 532.4 | 13.5 | 998.3 | 1011.4 | 13.1 |
| 531.4 | 544.9 | 13.5 | 1010.4 | 1021.9 | 11.5 |
| 543.9 | 558.0 | 14.1 | 1020.9 | 1031.1 | 10.2 |
| 557.0 | 571.1 | 14.1 | 1030.1 | 1041.6 | 11.5 |
| 570.1 | 583.6 | 13.5 | 1040.6 | 1054.6 | 14.0 |
| 582.6 | 598.0 | 15.4 | 1053.6 | 1067.6 | 14.0 |
| 597.0 | 614.3 | 17.3 | 1066.6 | 1081.5 | 14.9 |
| 613.3 | 630.5 | 17.2 | 1080.5 | 1091.5 | 11.0 |
| 629.5 | 646.8 | 17.3 | 1090.5 | 1100.8 | 10.3 |
| 645.8 | 662.4 | 16.6 | 1099.8 | 1107.9 | 8.1 |
| 661.4 | 679.3 | 17.9 | 1106.9 | 1114.6 | 7.7 |
| 678.3 | 696.8 | 18.5 | 1113.6 | 1120.5 | 6.9 |
| 695.8 | 714.9 | 19.1 | 1119.5 | 1126.4 | 6.9 |
| 713.9 | 733.6 | 19.7 | 1125.4 | 1132.2 | 6.8 |
| 732.6 | 753.6 | 21.0 | 1131.2 | 1138.1 | 6.9 |
| 752.6 | 774.3 | 21.7 | 1137.1 | 1143.6 | 6.5 |
| 773.3 | 795.5 | 22.2 | 1142.6 | 1149.0 | 6.4 |
| 794.5 | 817.4 | 22.9 | 1148.0 | 1155.3 | 7.3 |
| 816.4 | 841.8 | 25.4 | 1154.3 | 1161.2 | 6.9 |
| 840.8 | 868.0 | 27.2 | 1160.2 | 1167.9 | 7.7 |
| 867.0 | 899.3 | 32.3 | 1166.9 | 1175.4 | 8.5 |
| 898.3 | 933.0 | 34.7 | 1174.4 | 1184.3 | 9.9 |
| 932.0 | 966.8 | 34.8 | 1183.3 | 1193.1 | 9.8 |
| 965.8 | 1007.4 | 41.6 | 1192.1 | 1200.6 | 8.5 |
| 1006.4 | 1054.9 | 48.5 | 1199.6 | 1211.1 | 11.5 |
| 1053.9 | 1103.6 | 49.7 | 1210.1 | 1222.4 | 12.3 |
| 1102.6 | 1161.1 | 58.5 | 1221.4 | 1235.0 | 13.6 |
| 1160.1 | 1247.4 | 87.3 | 1234.0 | 1245.1 | 11.1 |

### Glycopeptide optimized variable window SWATH improves DIA glycopeptide measurement

Yeast cell wall proteins are highly *N*-glycosylated, and this subcellular fraction has been well studied by MS glycoproteomics [34, 37, 56]. Moreover, these studies have established that yeast strains with mutations in genes encoding enzymes in the *N*-glycosylation pathway have defects in the occupancy and structure of *N*-glycans in their cell wall glycoproteins. In this study, we tested the performance of our glycopeptide-optimized variable window SWATH method in measuring the glycosylation status of cell wall glycoproteins from yeast cells with defects in the *N*-glycosylation biosynthetic machinery. Glycosyltransferases encoded by *ALG* genes are responsible for assembly of the lipid linked oligosaccharide (LLO) donor substrate for *N*-glycosylation. *ALG3* encodes the Alg3p α-1,3-mannosyltransferase, which catalyzes the attachment of the sixth mannose to the growing LLO. *alg3Δ* mutant yeast cells accumulate Man_5_GlcNAc_2_ [61], resulting in lower glycosylation occupancy and truncated *N*-glycan structures on cell wall glycoproteins [34]. We used three different SWATH methods, fixed window SWATH (hereafter fixSWATH), vwSWATH, and gpvwSWATH, to measure peptides and glycopeptides from cell wall proteins from wild type and *alg3Δ* yeast. Interrogation of DIA data requires a high-quality ion library from peptide identifications from DDA data or generated theoretically. We used a previously published manually curated ion library of glycopeptides from yeast cell wall proteins [34].

The clear advantage of the narrower windows of gpvwSWATH in the *m/z* range rich in glycopeptides was apparent when inspecting MS1 spectra. For example, in the retention time range over which differently glycosylated forms of the Gas1-N^40^ (FFYSNN^40^GSQFYIR) glycopeptides eluted, glycopeptides were detected with Man_9_GlcNAc_2_ attached at an *m/z* of 1169.80^3+^ and with Man_10_GlcNAc_2_ attached at an *m/z* of 1223.82^3+^. Both of these glycoforms fell within the same vwSWATH window, because of the wide windows at this *m/z* range in this method optimized for unmodified peptides (Fig. 2A). These glycopeptides were also not resolved by LC retention time. It was therefore not possible to distinguish these glycopeptides by vwSWATH, although the overall signal intensity measured by this method was high because of the summed signal of the two glycopeptide forms (Fig. 2B). Also consistent with this lack of glycoform resolution, the oxonium ion intensities from vwSWATH were more than double that measured by the other two methods. The 25 *m/z* window in the fixSWATH method was theoretically small enough to distinguish glycan structures on the same peptide. However, the window still contained signals from other co-eluting peptides, and narrower windows will therefore improve specificity. This would be particularly relevant for mammalian glycans, where mass differences between glycoforms can be much smaller than in high mannose yeast glycans.

**Figure 2.**
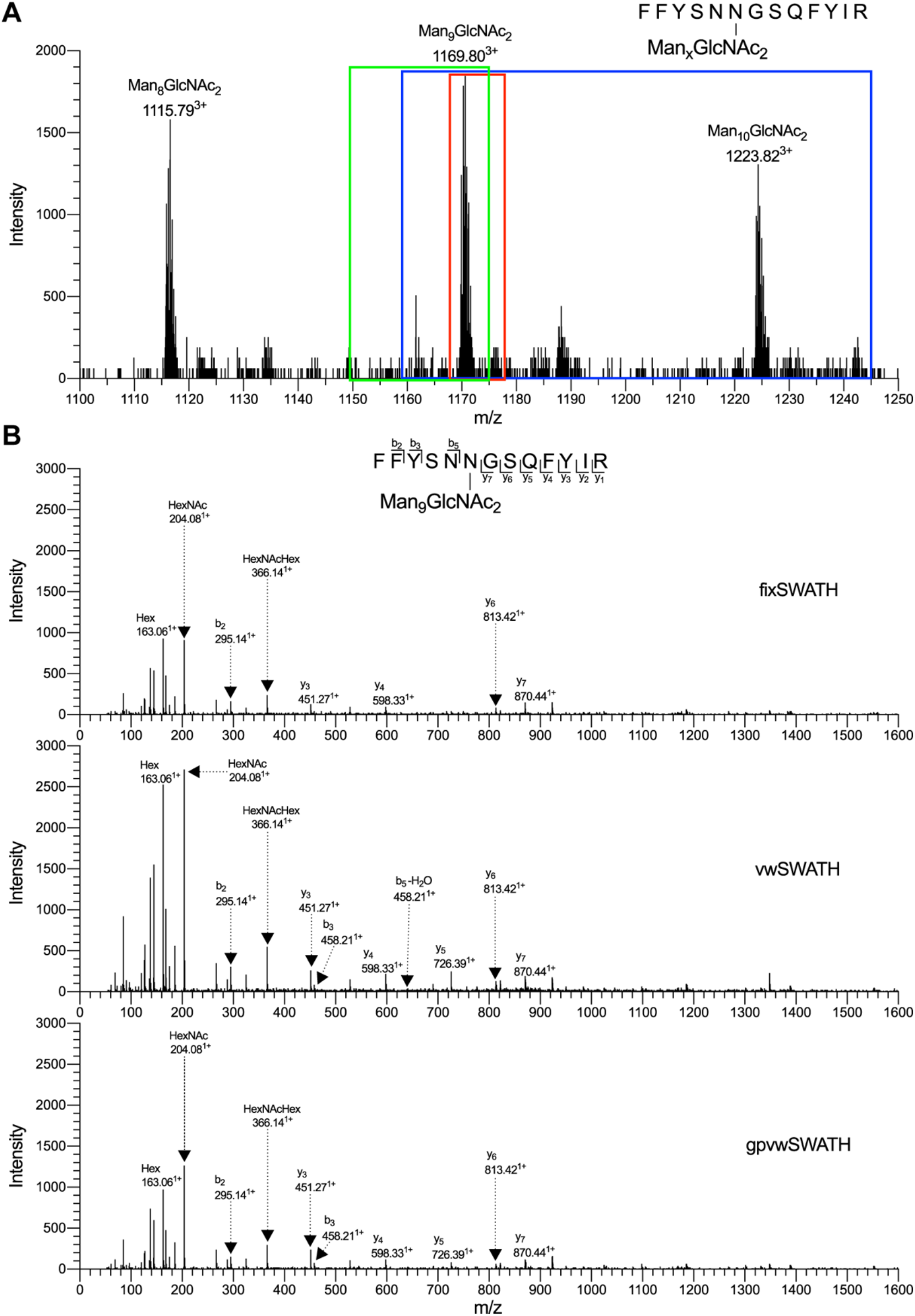
gpvwSWATH provides small windows for accurate glycopeptide measurement. **(A)** MS spectrum containing *N*-glycopeptide Gas1-N^40^ (FFYSNN^40^GSQFYIR) with Man_8_GlcNAc_2_, Man_9_GlcNAc_2_, and Man_10_GlcNAc_2_ attached. Windows provided by different SWATH methods are shown by colored frames: green, fixSWATH; blue, vwSWATH; red, gpvwSWATH. **(B)** MS/MS spectra measured by each SWATH method.

In *alg3*Δ yeast, glycan biosynthesis stops after five mannoses are added to the LLO, leading to accumulation of the Man_5_GlcNAc_2_ LLO [15]. This incomplete glycan is still transferred to protein substrates and can be further modified, forming various glycan structures on mature glycoproteins [15]. We tested the ability of gpvwSWATH, vwSWATH, and fixSWATH to distinguish site-specific glycan structures in *alg3Δ* and wild type yeast, focussing on analysis of the robustly measurable Gas1-N^40^ and Gas1-N^253^ glycosylation sites. Previous work has established that in *alg3*Δ yeast, Gas1-N^40^ is mostly modified with Man_5_GlcNAc_2_, with some Man_6_GlcNAc_2_ and Man_7_GlcNAc_2_ also present [34]. In contrast, Gas1-N^40^ in wild type yeast is glycosylated with high occupancy, and mostly carries the Man_9_GlcNAc_2_ glycan structure, as well as some Man_10_GlcNAc_2_ and Man_8_GlcNAc_2_ [34]. We found that all three SWATH methods successfully established that *alg3*Δ yeast had predominantly Man_5_GlcNAc_2_, and wild type yeast mainly Man_9_GlcNAc_2_ (Fig. 3A). However, the apparent intensities of Man_9_GlcNAc_2_ and Man_10_GlcNAc_2_ glycans at Gas1-N^40^ as measured by vwSWATH were identical (Fig. 3A). This incorrect assignment was due to both of these glycopeptides falling within the same wide window in vwSWATH (Fig. 2A). Statistical comparison of wild type and *alg3*Δ yeast identified significant differences in the intensities of Gas1-N^40^ glycopeptides containing 8, 9, and 10 mannoses when measured by fixSWATH or gpvwSWATH (Fig. 3B). However, fixSWATH and vwSWATH were able to detect the significant difference in abundance of the Man_5_GlcNAc_2_ glycopeptide, while gpvwSWATH could detect the difference in the Man_7_GlcNAc_2_ glycopeptide (Fig. 3B). The failure of gpvwSWATH to detect the difference in abundance of the Gas1-N^40^ Man_5_GlcNAc_2_ glycopeptide is likely because its small size places it within a wide *m/z* window in this method. This highlights that any given SWATH window design will not be optimal for all analytes of interest – in this case glycopeptides with relatively low *m/z* values. Inspection of glycosylation at Gas1-N^253^ showed similar performance of the three methods. Wild type yeast has mostly Man_11_GlcNAc_2_ on Gas1-N^253^, followed by Man_10_GlcNAc_2_ and Man_12_GlcNAc_2_, while *alg3*Δ yeast accumulates Man_7_GlcNAc_2_ and Man_8_GlcNAc_2_ [34]. The performance of vwSWATH in measuring Gas1-N^253^ glycopeptides was poor, as no glycopeptide carrying more than 10 mannoses was able to be clearly distinguished. Furthermore, Man_7_GlcNAc_2_, the dominant glycan structure in *alg3*Δ yeast, was not confidently detected at all by vwSWATH (Fig. 3C). Although the most intense glycan structures were accurately measured by both fixSWATH and gpvwSWATH, fixSWATH failed to detect several glycopeptides with low intensity, such as Man_9_GlcNAc_2_ and Man_10_GlcNAc_2_ in *alg3*Δ yeast (Fig. 3C). The failure of glycopeptides to be measured in these various samples was primarily due to difficulties in accurate automated peak picking in the more complex samples associated with wider windows. Finally, more glycopeptide forms were found to have significant differences in abundance between wild type and *alg3*Δ yeast when measured with gpvwSWATH than by fixSWATH, while no significant differences were detected using vwSWATH (Fig. 3D). In summary, we found that gpvwSWATH was the best approach for accurate and robust detection and measurement of glycopeptides, especially those with large glycans.

**Figure 3.**
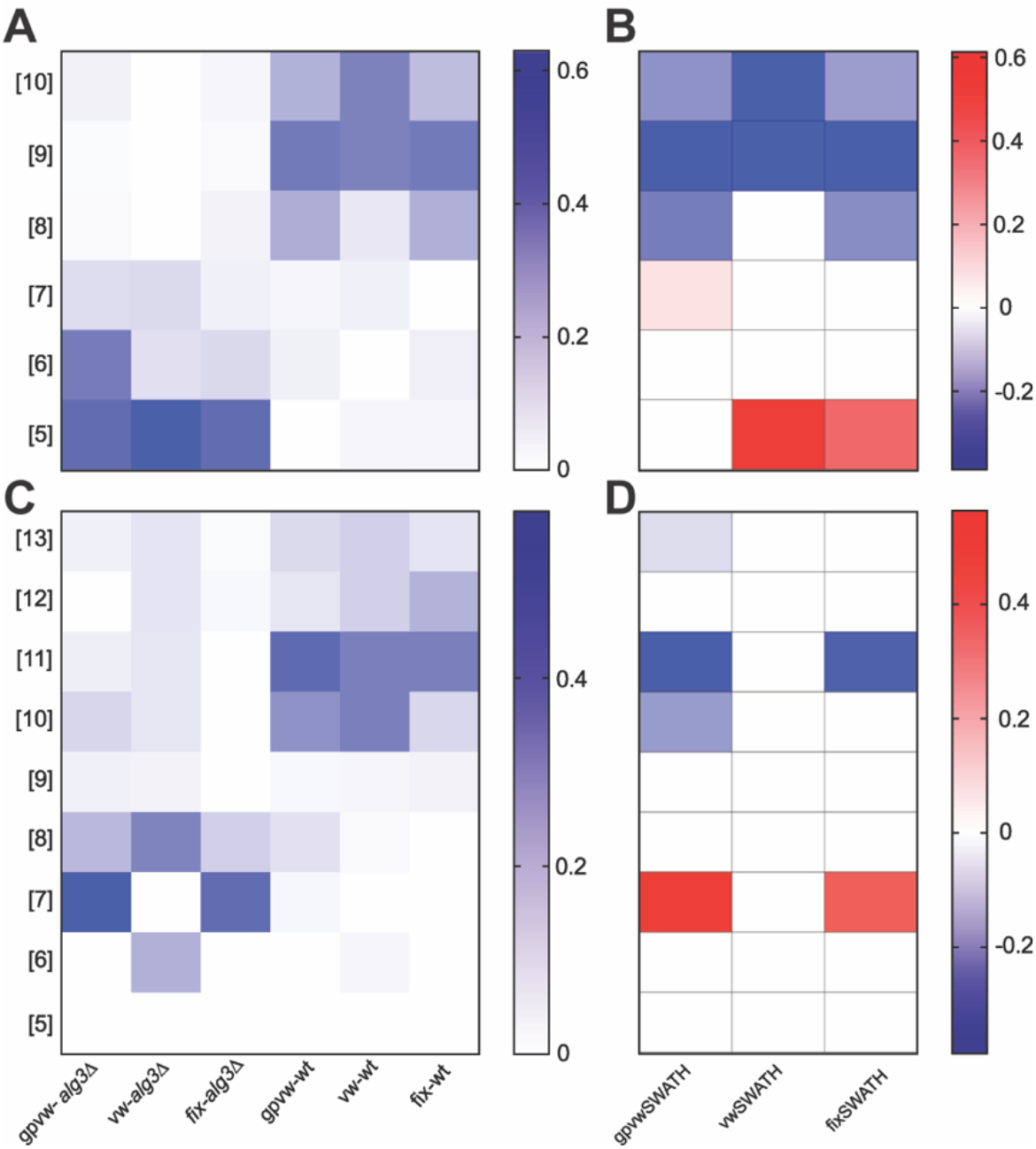
gpvwSWATH allows accurate measurement of glycopeptide abundance. Heatmap depicting intensities of **(A)** Gas1-N^40^ and **(C)** Gas-N^253^ glycopeptides with *N*-glycans containing two GlcNAc and an increasing number of mannoses in wild type and *alg3Δ* yeast cell wall samples, measured by the three SWATH methods. Heatmap showing the differences of the intensities of **(B)** Gas-N^40^ and **(D)** Gas-N^253^ glycopeptides with different glycoforms between *alg3Δ* and wild type yeast cell wall samples, measured by the three SWATH methods. White, not significant; red, higher in *alg3Δ*; blue, higher in wild type.

## Discussion

Improved analytical methods for detecting and measuring the abundance of glycopeptides are critical for efficient investigation of the many biological and industrial roles of protein glycosylation. DIA/SWATH glycoproteomics has been shown to be a powerful approach for analysing site-specific glycosylation in many diverse applications [34, 50, 62, 63]. The original fixed window width implementation of SWATH was sufficient for general peptide measurement [46], but could be improved with variable window SWATH, which involved decreasing the window size at precursor-ion dense regions of the *m/z* range to increase signal:noise and measure spectra with reduced complexity [51, 52]. However, with their large covalent oligosaccharide modifications, glycopeptides have a distinct *m/z* distribution to nonmodified peptides (Fig. 1). We therefore designed a variable window SWATH method optimized for detection and quantification of glycopeptides, and showed that it indeed demonstrated superior analytical performance (Fig. 2 and 3).

To generate a set of variable SWATH windows optimized for glycopeptide analysis we used the SWATH variable window calculator with a list of precursor masses whose MS/MS fragmentation spectra contained oxonium ions, identifying the precursor as a glycopeptide. We obtained these data from DDA analyses of wild type and glycosylation-deficient yeast cell wall samples. It would be possible to generate alternative variable window methods based on a precursor mass list generated from samples enriched in glycopeptides, from alternative glycosylation mutants, or alternative species, depending on the characteristics of the samples to be analysed. The gpvwSWATH method we used here showed superior analytical performance especially for large glycopeptides. However, some glycopeptides with smaller oligosaccharides were primarily detected with *m/z* precursor ions that fell in relatively wide windows, causing suboptimal measurement of these glycopeptides. While this is unfortunately an unavoidable general analytical feature of variable window SWATH methods, several alternative approaches are possible. Increasing the total number of windows will improve precursor specificity, although at the cost of reduced temporal resolution with an increased MS cycle time, potentially leading to less robust LC peak detection. Narrowing the overall *m/z* measurement range to exclude *m/z* regions with few target glycopeptides can also partially overcome these problems. DIA analysis of more complex glycoproteomes may be challenging, including of mammalian glycans with high structural heterogeneity. DIA relies on high quality ion libraries based on previous comprehensive identification of analytes of interest in a sample, and particular care should be taken in window design for glycopeptide analysis to ensure different structurally related glycopeptides of interest fall in separate windows. Even with very small *m/z* windows, structurally related glycopeptides may fall in the same SWATH window, and in this case accurate differentiation or measurement would not be possible. This limitation should be carefully considered when designing DIA methods and when interpreting glycopeptide-focused DIA data. The precise experimental question at hand will determine the optimal choice of method.

In conclusion, glycopeptide variable window SWATH is a sensible method for detection and quantification of glycopeptides with diverse glycan structures. Appropriate instrument methods can be quickly and easily generated and flexibly adjusted depending on the sample types, analytes of interest, and experimental priorities.

## Acknowledgements

We thank Dr Amanda Nouwens and Peter Josh at The University of Queensland, School of Chemistry and Molecular Biosciences Mass Spectrometry Facility for their assistance and expertise. BLS was funded by an Australian National Health and Medical Research Council RD Wright Biomedical (CDF Level 2) Fellowship APP1087975. This work was funded by an Australian Research Council Discovery Project DP160102766 and an Australian Research Council Industrial Transformation Training Centre IC160100027 to BLS.

## References

1. Spiro, R.G., Protein glycosylation: nature, distribution, enzymatic formation, and disease implications of glycopeptide bonds. Glycobiology, 2002. 12(4): p. 43R–56R.

2. Varki, A., Biological roles of glycans. Glycobiology, 2016. 27(1): p. 3–49.

3. Wormald, M.R. and R.A. Dwek, Glycoproteins: glycan presentation and protein-fold stability. Structure, 1999. 7(7): p. R155–R160.

4. Janik, M.E., A. Lityńska, and P. Vereecken, Cell migration-The role of integrin glycosylation. Biochimica et Biophysica Acta (BBA) - General Subjects, 2010. 1800(6): p. 545–555.

5. Haines, N. and K.D. Irvine, Glycosylation regulates Notch signalling. Nature Reviews Molecular Cell Biology, 2003. 4: p. 786–797.

6. Xu, C. and D.T. Ng, Glycosylation-directed quality control of protein folding. Nature Reviews Molecular Cell Biology, 2015. 16(12): p. 742–752.

7. Quast, I., B. Peschke, and J.D. Lünemann, Regulation of antibody effector functions through IgG Fc N-glycosylation. Cellular and Molecular Life Sciences, 2017. 74(5): p. 837–847.

8. Apweiler, R., H. Hermjakob, and N. Sharon, On the frequency of protein glycosylation, as deduced from analysis of the SWISS-PROT database11Dedicated to Prof. Biochimica et Biophysica Acta (BBA) - General Subjects, 1999. 1473(1): p. 4-8.

9. Zielinska, Dorota F., F. Gnad, K. Schropp, Jacek R. Wiśniewski, and M. Mann, Mapping N-glycosylation sites across seven evolutionarily distant species reveals a divergent substrate proteome despite a common core machinery. Molecular Cell, 2012. 46(4): p. 542–548.

10. Trombetta, E.S., The contribution of N-glycans and their processing in the endoplasmic reticulum to glycoprotein biosynthesis. Glycobiology, 2003. 13(9): p. 77R–91R.

11. Aebi, M., R. Bernasconi, S. Clerc, and M. Molinari, N-glycan structures: recognition and processing in the ER. Trends in Biochemical Sciences, 2010. 35(2): p. 74–82.

12. Xiao, H., J.M. Smeekens, and R. Wu, Quantification of tunicamycin-induced protein expression and N-glycosylation changes in yeast. Analyst, 2016. 141(12): p. 3737–3745.

13. Aebi, M., N-linked protein glycosylation in the ER. Biochimica et Biophysica Acta (BBA) - Molecular Cell Research, 2013. 1833(11): p. 2430–2437.

14. Schulz, B.L., Beyond the sequon: sites of N-glycosylation, in Glycosylation. 2012, IntechOpen.

15. Burda, P. and M. Aebi, The dolichol pathway of N-linked glycosylation. Biochimica et Biophysica Acta (BBA) - General Subjects, 1999. 1426(2): p. 239–257.

16. Poljak, K., N. Selevsek, E. Ngwa, J. Grossmann, M.E. Losfeld, and M. Aebi, Quantitative Profiling of N-linked Glycosylation Machinery in Yeast <Saccharomyces cerevisiae>. Molecular & Cellular Proteomics, 2018. 17(1): p. 18.

17. Roth, J., Protein N-Glycosylation along the secretory pathway: Relationship to organelle topography and function, protein quality control, and cell interactions. Chemical Reviews, 2002. 102(2): p. 285–304.

18. Arigoni-Affolter, I., E. Scibona, C.-W. Lin, D. Brühlmann, J. Souquet, H. Broly, and M. Aebi, Mechanistic reconstruction of glycoprotein secretion through monitoring of intracellular N-glycan processing. Science Advances, 2019. 5(11): p. eaax8930.

19. Arnold, J.N., R. Saldova, U.M.A. Hamid, and P.M. Rudd, Evaluation of the serum-N linked glycome for the diagnosis of cancer and chronic inflammation. Proteomics, 2008.8(16): p. 3284–3293.

20. Chandler, K. and R. Goldman, Glycoprotein disease markers and single proteinomics. Molecular & Cellular Proteomics, 2013. 12(4): p. 836–845.

21. Oliveira-Ferrer, L., K. Legler, and K. Milde-Langosch, Role of protein glycosylation in cancer metastasis. Seminars in Cancer Biology, 2017. 44: p. 141–152.

22. Thaysen-Andersen, M., N.H. Packer, and B.L. Schulz, Maturing glycoproteomics technologies provide unique structural insights into the N-glycoproteome and its regulation in health and disease. Molecular & Cellular Proteomics, 2016. 15(6): p. 1773–1790.

23. Jaeken, J. and G. Matthijs, Congenital disorders of glycosylation: a rapidly expanding disease family. Annu. Rev. Genomics Hum. Genet., 2007. 8: p. 261–278.

24. Freeze, H.H. and M. Aebi, Altered glycan structures: the molecular basis of congenital disorders of glycosylation. Current Opinion in Structural Biology, 2005. 15(5): p. 490–498.

25. Novak, J., B.A. Julian, J. Mestecky, and M.B. Renfrow, Glycosylation of IgA1 and pathogenesis of IgA nephropathy. Seminars in Immunopathology, 2012. 34(3): p. 365–382.

26. Narimatsu, H., H. Kaji, S.Y. Vakhrushev, H. Clausen, H. Zhang, E. Noro, A. Togayachi, C. Nagai-Okatani, A. Kuno, X. Zou, L. Cheng, S.-C. Tao, and Y. Sun, Current Technologies for Complex Glycoproteomics and Their Applications to Biology/Disease-Driven Glycoproteomics. Journal of Proteome Research, 2018. 17(12): p. 4097–4112.

27. Freeze, H.H., E.A. Eklund, B.G. Ng, and M.C. Patterson, Neurology of inherited glycosylation disorders. The Lancet. Neurology, 2012. 11(5): p. 453–466.

28. Peracaula, R., S. Barrabés, A. Sarrats, P.M. Rudd, and R. de Llorens, Altered glycosylation in tumours focused to cancer diagnosis. Disease Markers, 2008. 25(4-5): p. 207–218.

29. Reis, C.A., H. Osorio, L. Silva, C. Gomes, and L. David, Alterations in glycosylation as biomarkers for cancer detection. Journal of Clinical Pathology, 2010. 63(4): p. 322–329.

30. Adamczyk, B., T. Tharmalingam, and P.M. Rudd, Glycans as cancer biomarkers. Biochimica et Biophysica Acta (BBA) - General Subjects, 2012. 1820(9): p. 1347–1353.

31. Zacchi, L.F. and B.L. Schulz, N-glycoprotein macroheterogeneity: biological implications and proteomic characterization. Glycoconjugate Journal, 2016. 33(3): p. 359–376.

32. Dell, A. and H.R. Morris, Glycoprotein structure determination by mass spectrometry. Science, 2001. 291(5512): p. 2351–2356.

33. Ruhaak, L.R., G. Xu, Q. Li, E. Goonatilleke, and C.B. Lebrilla, Mass spectrometry approaches to glycomic and glycoproteomic analyses. Chemical Reviews, 2018. 118(17): p. 7886–7930.

34. Zacchi, L.F. and B.L. Schulz, SWATH-MS glycoproteomics reveals consequences of defects in the glycosylation machinery. Molecular & Cellular Proteomics, 2016. 15(7): p. 2435–2447.

35. Mayampurath, A., E. Song, A. Mathur, C.-y. Yu, Z. Hammoud, Y. Mechref, and H. Tang, Label-free glycopeptide quantification for biomarker discovery in human sera. Journal of proteome research, 2014. 13(11): p. 4821–4832.

36. Song, E., S. Pyreddy, and Y. Mechref, Quantification of glycopeptides by multiple reaction monitoring liquid chromatography/tandem mass spectrometry. Rapid Communications in Mass Spectrometry, 2012. 26(17): p. 1941–1954.

37. Xu, Y., U.M. Bailey, and B.L. Schulz, Automated measurement of site specific-N glycosylation occupancy with SWATH-MS. Proteomics, 2015. 15(13): p. 2177–2186.

38. Liu, Z., J. Cao, Y. He, L. Qiao, C. Xu, H. Lu, and P. Yang, Tandem 18O stable isotope labeling for quantification of N-glycoproteome. Journal of Proteome Research, 2009. 9(1): p. 227–236.

39. Sun, Z., H. Qin, F. Wang, K. Cheng, M. Dong, M. Ye, and H. Zou, Capture and dimethyl labeling of glycopeptides on hydrazide beads for quantitative glycoproteomics analysis. Analytical Chemistry, 2012. 84(20): p. 8452–8456.

40. Zhou, Y., Y. Shan, L. Zhang, and Y. Zhang, Recent advances in stable isotope labeling based techniques for proteome relative quantification. Journal of Chromatography A, 2014. 1365: p. 1–11.

41. Rebecchi, K.R., J.L. Wenke, E.P. Go, and H. Desaire, Label-free quantitation: a new glycoproteomics approach. Journal of the American Society for Mass Spectrometry, 2009.20(6): p. 1048–1059.

42. Dahabiyeh, L.A., D. Tooth, and D.A. Barrett, Profiling of 54 plasma glycoproteins by label-free targeted LC-MS/MS. Analytical Biochemistry, 2019. 567: p. 72–81.

43. Röst, H.L., G. Rosenberger, P. Navarro, L. Gillet, S.M. Miladinović, O.T. Schubert, W. Wolski, B.C. Collins, J. Malmström, and L. Malmström, OpenSWATH enables automated, targeted analysis of data-independent acquisition MS data. Nature Biotechnology, 2014. 32(3): p. 219–223.

44. Egertson, J.D., A. Kuehn, G.E. Merrihew, N.W. Bateman, B.X. MacLean, Y.S. Ting, J.D. Canterbury, D.M. Marsh, M. Kellmann, and V. Zabrouskov, Multiplexed MS/MS for improved data-independent acquisition. Nature Methods, 2013. 10(8): p. 744–746.

45. Lange, V., P. Picotti, B. Domon, and R. Aebersold, Selected reaction monitoring for quantitative proteomics: a tutorial. Molecular Systems Biology, 2008. 4(1): p. 222.

46. Gillet, L.C., P. Navarro, S. Tate, H. Röst, N. Selevsek, L. Reiter, R. Bonner, and R. Aebersold, Targeted data extraction of the MS/MS spectra generated by data-independent acquisition: a new concept for consistent and accurate proteome analysis. Molecular & Cellular Proteomics, 2012. 11(6): p. O111–016717.

47. Wu, W., W.W. Yong, and M.C.M. Chung, A simple biomarker scoring matrix for early gastric cancer detection. Proteomics, 2016. 16(22): p. 2921–2930.

48. Gajbhiye, A., R. Dabhi, K. Taunk, G. Vannuruswamy, S. RoyChoudhury, R. Adhav, S. Seal, A. Mane, S. Bayatigeri, and M.K. Santra, Urinary proteome alterations in HER2 enriched breast cancer revealed by multipronged quantitative proteomics. Proteomics, 2016. 16(17): p. 2403–2418.

49. Liu, Y., J. Chen, A. Sethi, Q.K. Li, L. Chen, B. Collins, L.C. Gillet, B. Wollscheid, H. Zhang, and R. Aebersold, Glycoproteomic analysis of prostate cancer tissues by SWATH mass spectrometry discovers N-acylethanolamine acid amidase and protein tyrosine kinase 7 as signatures for tumor aggressiveness. Molecular & Cellular Proteomics, 2014. 13(7): p. 1753–1768.

50. Liu, T., S. Shang, W. Li, X. Qin, L. Sun, S. Zhang, and Y. Liu, Assessment of hepatocellular carcinoma metastasis glycobiomarkers using advanced quantitative N-glycoproteome analysis. Frontiers in Physiology, 2017. 8: p. 472.

51. Hunter, C. and S. Seymour, Improved data quality using variable Q1 window widths in SWATH™ acquisition-Data independent acquisition on the TripleTOF^®^ 6600 and 5600+ systems. AB SCIEX Technical Note, 2015: p. 10090114–01.

52. Zhang, Y., A. Bilbao, T. Bruderer, J. Luban, C. Strambio-De-Castillia, F. Lisacek, G. Hopfgartner, and E. Varesio, The use of variable Q1 isolation windows improves selectivity in LC-SWATH-MS acquisition. Journal of Proteome Research, 2015. 14(10): p. 4359–4371.

53. Bruderer, R., O.M. Bernhardt, T. Gandhi, S.M. Miladinović, L.-Y. Cheng, S. Messner, T. Ehrenberger, V. Zanotelli, Y. Butscheid, and C. Escher, Extending the limits of quantitative proteome profiling with data-independent acquisition and application to acetaminophen-treated three-dimensional liver microtissues. Molecular & Cellular Proteomics, 2015. 14(5): p. 1400–1410.

54. Meyer, J.G., A.K. D’Souza, D.J. Sorensen, M.J. Rardin, A.J. Wolfe, B.W. Gibson, and B. Schilling, Quantification of lysine acetylation and succinylation stoichiometry in proteins using mass spectrometric data-independent acquisitions (SWATH). Journal of the American Society for Mass Spectrometry, 2016. 27(11): p. 1758–1771.

55. Sidoli, S., S. Lin, L. Xiong, N.V. Bhanu, K.R. Karch, E. Johansen, C. Hunter, S. Mollah, and B.A. Garcia, Sequential window acquisition of all theoretical mass spectra (SWATH) analysis for characterization and quantification of histone post-translational modifications. Molecular & Cellular Proteomics, 2015. 14(9): p. 2420–2428.

56. Schulz, B.L. and M. Aebi, Analysis of glycosylation site occupancy reveals a role for Ost3p and Ost6p in site-specific N-glycosylation efficiency. Molecular & Cellular Proteomics, 2009. 8(2): p. 357–364.

57. Phung, T.K., L.F. Zacchi, and B.L. Schulz, DIALib: an automated ion library generator for data independent acquisition mass spectrometry analysis of peptides and glycopeptides. bioRxiv, 2019: p. 696518.

58. Kerr, E.D., C.H. Caboche, and B.L. Schulz, Post-translational modifications drive protein stability to control the dynamic beer brewing proteome. Molecular & Cellular Proteomics, 2019: p. mcp.RA119.001526.

59. Perez-Riverol, Y., A. Csordas, J. Bai, M. Bernal-Llinares, S. Hewapathirana, D.J. Kundu, A. Inuganti, J. Griss, G. Mayer, M. Eisenacher, E. Pérez, J. Uszkoreit, J. Pfeuffer, T. Sachsenberg, S. Yilmaz, S. Tiwary, J. Cox, E. Audain, M. Walzer, A.F. Jarnuczak, T. Ternent, A. Brazma, and J.A. Vizcaíno, The PRIDE database and related tools and resources in 2019: improving support for quantification data. Nucleic acids research, 2019. 47(D1): p. D442–D450.

60. Kerr, E.D., T.K. Phung, C.H. Caboche, G.P. Fox, G.J. Platz, and B.L. Schulz, The intrinsic and regulated proteomes of barley seeds in response to fungal infection. Analytical Biochemistry, 2019. 580: p. 30–35.

61. Huffaker, T.C. and P.W. Robbins, Yeast mutants deficient in protein glycosylation. Proceedings of the National Academy of Sciences, 1983. 80(24): p. 7466–7470.

62. Sanda, M., L. Zhang, N.J. Edwards, and R. Goldman, Site-specific analysis of changes in the glycosylation of proteins in liver cirrhosis using data-independent workflow with soft fragmentation. Analytical and Bioanalytical Chemistry, 2017. 409(2): p. 619–627.

63. Keller, A., S.L. Bader, U. Kusebauch, D. Shteynberg, L. Hood, and R.L. Moritz, Opening a SWATH window on posttranslational modifications: automated pursuit of modified peptides. Molecular & Cellular Proteomics, 2016. 15(3): p. 1151–1163.

